# The dark side of fluorescent protein tagging – the impact of protein tags on biomolecular condensation

**DOI:** 10.1101/2024.11.23.624970

**Authors:** Edoardo Fatti, Sarah Khawaja, Karsten Weis

## Abstract

Biomolecular condensation has emerged as an important mechanism to control various cellular processes through the formation of membraneless organelles. Fluorescent protein tags have been extensively used to study the formation and the properties of condensates *in vitro* and *in vivo*, but there is evidence that tags may perturb the condensation properties of proteins. In this study, we carefully assess the effects of protein tags on the yeast DEAD-box ATPase Dhh1, a central regulator of processing bodies (P-bodies), which are biomolecular condensates involved in mRNA metabolism. We show that fluorescent tags as well as a poly-histidine tag greatly affect Dhh1 condensation *in vitro* and lead to condensates with different dynamic properties. Tagging of Dhh1 with various fluorescent proteins *in vivo* alters the number of P-bodies upon glucose starvation and some tags even show constitutive P-bodies in non-stressed cells. These data raise concerns about the accuracy of tagged protein condensation experiments, highlighting the need for caution when interpreting the results.

**Significance Statement:**

- Fluorescent tags are extensively used in protein condensation studies although their effect in condensate dynamics has not been carefully investigated.
- Tags affect the condensation propensity and dynamics of Dhh1 *in vitro* and P-body numbers *in vivo*.
- Tags may generally alter the behavior of proteins in biomolecular condensates and their use needs to be carefully evaluated and controlled.

## Introduction

Protein condensation has emerged as a general mechanism that governs the formation of membraneless organelles, and is implicated in a growing number of processes in eukaryotic cells (Hyman *et al*., 2014; Banani *et al*., 2017; Shin and Brangwynne, 2017; Hirose *et al*., 2023). These condensates are promoted by weak, multivalent protein-protein and/or protein-RNA interactions, often involving low complexity domains (LCDs) (Choi *et al*., 2020; Hondele *et al*., 2020; Roden and Gladfelter, 2021). The plethora of eukaryotic membraneless organelles includes the nucleolus, nuclear speckles, Cajal bodies, processing bodies (P-bodies) and stress granules (Hyman *et al*., 2014; Banani *et al*., 2017; Shin and Brangwynne, 2017; Hirose *et al*., 2023). Condensates can differ greatly in their material properties, which are influenced by factors such as pH, temperature, salt concentration and composition. They can assemble and disassemble in response to changing environmental conditions, and in some cases have been shown to mature into irreversible aggregates over time and thus are thought to play a role in neurodegenerative aggregation disorders (Aguzzi and Altmeyer, 2016; Alberti and Dormann, 2019).

P-bodies are an example of dynamically remodeled protein condensates, and in yeast were identified as cytoplasmic foci that become microscopically visible following stress-induced translation repression (Parker and Sheth, 2007). Although their function is not fully understood, they are believed to be sites of mRNA storage and/or decay, since they are enriched for translation repressors and mRNA degradation factors such as Dhh1, Pat1, Xrn1 and Dcp1/2 (Sheth and Parker, 2003; Teixeira and Parker, 2007; Aizer *et al*., 2014; Mugler *et al*., 2016; Luo *et al*., 2018). The DEAD-box ATPase Dhh1/DDX6 has been identified as a critical component of P-bodies and is a key regulator of their dynamics in yeast. For example, inhibiting Dhh1’s ATPase function was shown to induce the formation of constitutive P-bodies even in the absence of stress, and deletion of Dhh1 dramatically reduces the number of P-bodies that form in stress (Carroll *et al*., 2011; Mugler *et al*., 2016).

The discovery of fluorescent protein (FP) variants has revolutionized our ability to observe the localization and dynamics of proteins in living cells and organisms (Snapp, 2009; Cranfill *et al*., 2016; Lambert, 2019) Countless studies have therefore made use of FP tags such as Green Fluorescent Protein (GFP) and mCherry to visualize membraneless organelles and to study components of biomolecular condensates. While *in vitro* phase contrast microscopy can be used to observe protein condensates, membraneless organelles in living cells need to be visualized with fluorescent tags, since their density/refractive index is similar to that of the intracellular space

(Schlüßler *et al*., 2022). Antibodies can be used in fixed cells, but fixation can cause artifacts and prevents any dynamic studies or assessment of material properties (Irgen-Gioro *et al*., 2022). Thus, FPs have been invaluable tools for observing and visualizing condensates, and indeed for studying practically all aspects of cell biology. The utility of FPs as reporters depends on their not affecting the normal behavior of the protein they are fused to. However, some recent studies have raised concerns that the use of these tags may alter the condensation properties of proteins (Feric *et al*., 2016; Uebel and Phillips, 2019; Barkley *et al*., 2024; Pandey *et al*., 2024). Unlike ‘classical’ protein complexes, condensates often rely on weak, multivalent interactions, and subtle changes in valency or interaction strength can have a large effect on the condensation behavior and the dynamics of condensates.

By extensively characterizing the phase separation properties of Dhh1 in a range of conditions, we show here that the addition of different protein tags greatly influences Dhh1 phase separation *in vitro* and *in vivo*. The addition of GFP or mCherry2 inhibits Dhh1 condensation but the addition of a hexahistidine (His) tag can oppose this effect. Condensate dynamics are also influenced by tags, producing different results depending on the tag used. Furthermore, the impact of tags is prominent *in vivo*, where both the type and the position of the tag affects protein expression levels and number of P-bodies, even promoting constitutive P-bodies in unstressed yeast cells. While we focus here on Dhh1, our results raise general concerns about the use of protein tags in the context of protein condensation and highlight the need to carefully interpret and control *in vitro* and *in vivo* experiments with protein tags.

## Results and discussion

### Tagging of Dhh1 affects its condensation *in vitro*

To examine how tags affect Dhh1 condensation, we carefully analyzed the influence of commonly used protein tags. We expressed and purified Dhh1 from *E. coli*, both untagged and tagged at the N-terminus with monomeric enhanced Green Fluorescent Protein (GFP) or mCherry2 (mCh2). Since histidine (His) tags are used extensively for biochemical purification and are often not removed before performing assays, we also included a His-tagged version of these protein constructs (Supplemental Figure S1A). We then performed *in vitro* protein condensation assays by mixing various concentrations of tagged or untagged Dhh1 with the RNA substrate polyuridylic acid (pU) and ATP at pH 6.0. We found that while untagged Dhh1 formed condensates starting at 1 μM, GFP-Dhh1 and mCh2-Dhh1 required 2 and 5 μM, respectively, to show condensates (Figure 1A, left panel). A His tag alone had no observable effect on Dhh1 condensation at any of the concentrations tested (Figure 1A), but it reversed the negative effect of GFP and mCh2 on condensation in double-tagged constructs (Figure 1A, right panel). In parallel, we tested if formation of Dhh1 condensates was dependent on RNA by omitting pU from the reaction mix. Untagged Dhh1 did not form condensates at any concentration but we observed condensates of His-Dhh1 and His-GFP-Dhh1 at the highest tested concentration (10 μM) (Supplemental Figure S2A, right panel). Surprisingly, 10 μM GFP-Dhh1 also formed condensates in the absence of pU (Supplemental Figure S2A, left panel). These results suggest that tagging of Dhh1 can have different effects on its ability to form condensates, with GFP and mCh2 overall increasing the critical concentration needed for condensate formation and the His tag reducing it.

**Figure 1.**
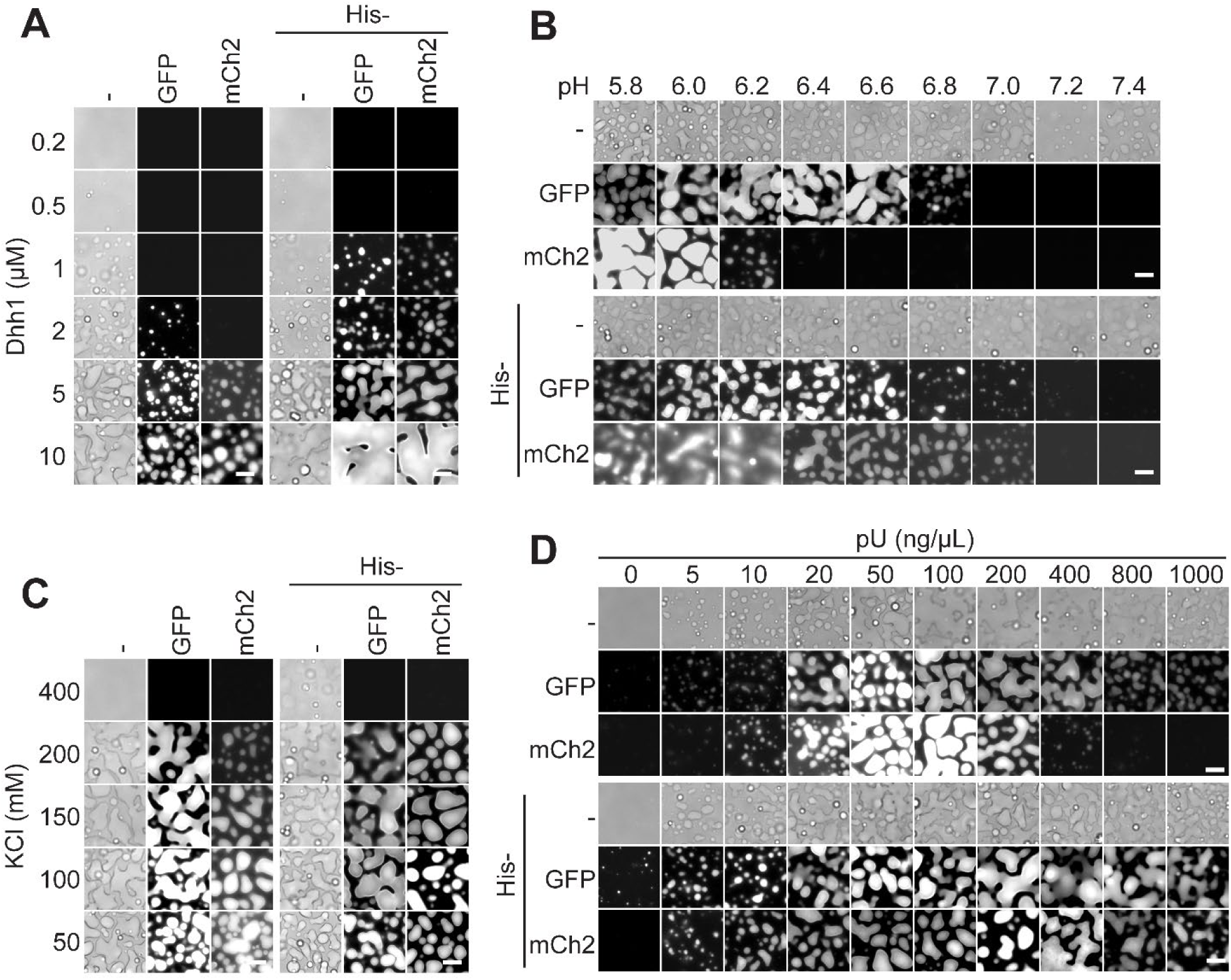
Tagging affects the condensation behavior of Dhh1 in the presence of RNA. (A) Left panel: Dhh1, GFP-Dhh1, and mCh2-Dhh1 form condensates at different protein concentrations. Right panel: His tag promotes GFP-Dhh1 and mCh2-Dhh1 condensation at lower concentrations. Assay performed in presence of pU, 100mM KCl and pH 6.0. (B) GFP and mCh2 tagging reduce Dhh1 condensation propensity at lower pHs in the presence of pU but a His tag reverts the effect. (C) Tagging of Dhh1 does not affect salt requirement in the presence of pU at pH 6.0. (D) GFP-Dhh1 and mCh2-Dhh1 show a lower propensity to form condensates at increasing pU concentrations at pH 6.0 compared to Dhh1 but a His tag reverts this effect. 5 mM ATP is present in all assays. Scale bar = 10 μm.

We previously showed that pH can influence the condensation behavior of Dhh1 (Hondele *et al*., 2019). To explore the effect of the tags in different pH conditions, we carried out a pH screen from pH 5.8 to 7.4 in the presence of RNA and ATP using 5 μM protein, a concentration where we saw a strict RNA-dependence on condensate formation for all constructs (Supplemental Figure S2A). Dhh1 formed condensates across the range of pHs, while GFP-Dhh1 only formed condensates up to pH 6.8 and mCh2-Dhh1 only up to pH 6.0 (Figure 1B, upper panel). Dhh1 and His-Dhh1 did not show any major differences, but the doubly tagged His-GFP-Dhh1 and His-mCh2-Dhh1 again showed a greater propensity to phase separate compared to the non-His-tagged versions, as condensates could be seen up to pH 7.0 in both cases (Figure 1B, lower panel). In the absence of pU, no condensates formed at any of the tested pHs with the protein concentration used (Supplemental Figure S2B).

Next, we looked at the effect of salt concentration, since ionic strength can modulate both protein-protein and protein-RNA interactions and therefore is also an important parameter for protein condensation. We found that titration of KCl did not significantly affect the condensation of Dhh1 in a tag-dependent manner, with condensation seen up to 200 mM KCl for all variants except His-Dhh1, which showed condensates even at 400 mM KCl (Figure 1C). At the lowest KCl concentration tested, GFP-Dhh1, His-Dhh1, His-GFP-Dhh1 and His-mCh2-Dhh1 formed condensates even in absence of pU (Supplemental Figure S2C).

Finally, we investigated the effect of varying pU concentration on the condensation of tagged and untagged Dhh1. Both Dhh1 and GFP-Dhh1 formed condensates from 5 ng/μL pU, the lowest non-zero concentration tested, and up to 1000 ng/μL, the highest tested concentration (Figure 1D, upper panel). Condensate size increased with increasing RNA concentration before plateauing and then decreasing again, at 800 ng/μL for Dhh1 and 400 ng/μL for GFP-Dhh1. mCh2-Dhh1 formed small condensates from 10 ng/μL pU, which increased in size until 100 ng/μL before becoming smaller and disappearing at 800 ng/μL. Addition of a His tag to all three constructs resulted in a similar phenotype to untagged Dhh1 across the range of pU concentrations, with the exception that small condensates could be seen with His-GFP-Dhh1 even in the absence of pU (Figure 1D, lower panel). These results are consistent with our observations that GFP and mCh2 inhibit Dhh1 condensation *in vitro* while the His tag reverts this effect and possibly even promotes condensation further.

### Spike-ins accurately report on condensate formation but not dynamics

Given the observed effects of protein tags, we wondered if we could develop a strategy to minimize the impact of the tag and provide unbiased information on the condensation properties of Dhh1. Our approach was to test if spiking in small amounts of GFP-Dhh1 in an otherwise unlabeled sample would be sufficient to visualize condensates without affecting their properties. First, we determined the minimal amount of GFP-Dhh1 to spike into Dhh1 condensates and found that both 1% and 5% give a satisfactory fluorescent signal, and we thus used 2% for further experiments (Figure 2A). We tested this ratio at a range of pH values and pU concentrations using GFP-Dhh1, mCh2-Dhh1 and, in parallel, a variant of Dhh1 labelled at the N-terminus with Cyanine 5 (Cy5-Dhh1), a small dye of 829 Da. In all cases tested, the spiked-in tagged variants did not alter the condensation propensity of untagged Dhh1 and did not affect the pH or pU requirements for condensation (Figure 2A, B and C). Furthermore, in all cases, the spiked-in protein completely co-localized with Dhh1 condensates (Figure 2B and C).

**Figure 2.**
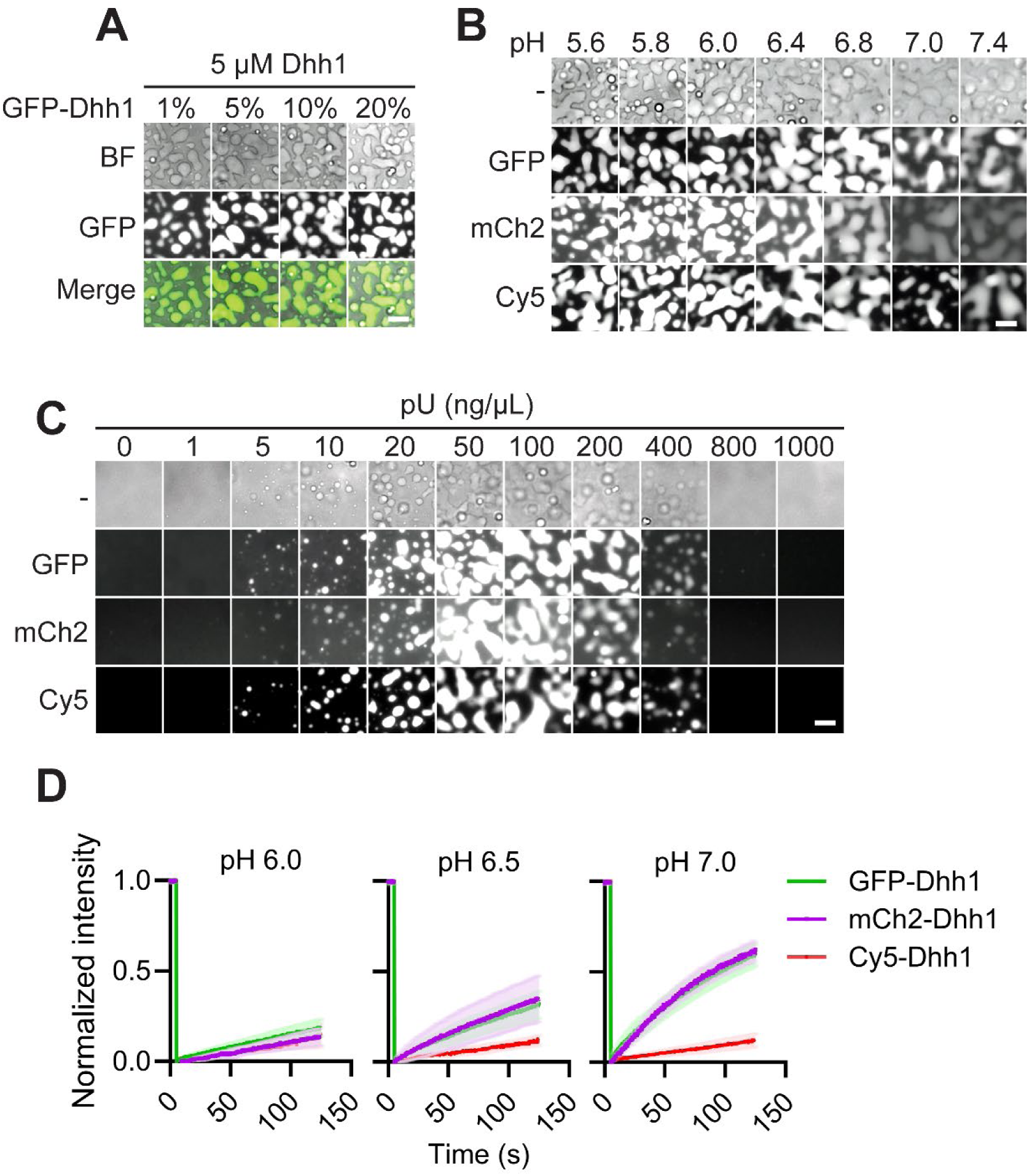
Spiking-in of tagged Dhh1 results in variable condensate dynamics as measured by FRAP (A) Different fractions of spiked-in GFP-Dhh1 co-localize with 5 μM Dhh1 in the presence of pU at pH 6.0. (B) Spiking-in of 2 % GFP-, mCh2-, and Cy5-Dhh1 with 5 μM Dhh1 in the presence of pU does not affect the pH dependence of Dhh1 condensation. (C) Spiking-in of 2 % GFP-, mCh2-, and Cy5-Dhh1 with 5 μM Dhh1 at pH 6.0 does not affect Dhh1 condensation at different pU concentrations. (D) Spiking-in of 2 % GFP -, mCh2-, and Cy5-Dhh1 with 5 μM Dhh1 in the presence of pU at different pHs shows different FRAP recovery rates. N = 2. 100mM KCl and 5 mM ATP are present in all assays. Scale bar = 10 μm.

To test if spike-ins can report on the dynamic exchange of Dhh1 into and out of condensates, we performed FRAP measurements on *in vitro*-formed droplets at different pHs. At pH 6.0, all three tagged proteins showed similar recovery rates, with GFP-Dhh1 slightly faster than mCh2-Dhh1 and Cy5-Dhh1 (Figure 2D). However, at pH 6.5 and more so at pH 7.0, GFP-Dhh1 and mCh2-Dhh1 had a significantly faster fluorescence recovery rate than Cy5-Dhh1 (Figure 2D), despite showing a similar propensity to phase separate (Figure 2B). Doubly tagged His-GFP-Dhh1 and His-mCh2-Dhh1 also showed a similar trend, with increasing recovery rates at higher pH but overall lower rates compared to GFP-Dhh1 and mCh2-Dhh1 (Supplemental Figure S3A and B).

We conclude that a low ratio of spiked-in protein variants does not seem to interfere with the critical concentration for bulk condensation of Dhh1 and spike-ins can thus be a valuable tool to study condensation formation under different conditions. However, spike-ins can still not be readily used to assess condensate dynamics by FRAP. The behavior of the bulk, untagged protein remains invisible in this assay and only the dynamics of the small fraction of the tagged protein is revealed, which is likely still influenced by the tag, e.g., by affecting its interaction properties within the condensate. Furthermore, these data also show that buffer conditions, such as pH, can have a strong influence on the recovery rate and FRAP should be ideally performed at close physiological buffer conditions when possible.

### Fluorescent tags affect P-body numbers *in vivo*

The drastic effects of tags on Dhh1 condensation *in vitro* raise questions about their impact *in vivo*. Since Dhh1 has been shown to be critical for the formation of P-bodies in budding yeast upon stress (Mugler *et al*., 2016), we examined the effect of tagging Dhh1 on the formation of P-bodies upon acute glucose deprivation. To recapitulate our *in vitro* experiments, we first tagged endogenous Dhh1 at the N-terminus with GFP, His-GFP, mCh2 and His-mCh2 and imaged cells in glucose-rich (+gluc) and glucose-deplete (-gluc) medium. Consistent with previous results, P-bodies were not detectable in any of the strains in +gluc, (Figure 3A). In -gluc, GFP-Dhh1 and His-GFP-Dhh1 showed a comparable number of P-bodies, with a slight increase for His-GFP-Dhh1. By contrast, mCh2-Dhh1 and His-mCh2-Dhh1 showed approximately 40-50% fewer P-bodies compared to GFP-Dhh1, again with slightly higher P-body numbers for the His-tagged variant (Figure 3A and B). This result is consistent with our *in vitro* data showing that mCh2 acts as a stronger solubilizer than GFP and that the His tag promotes Dhh1 condensation. Notably, western blot analysis showed that tags also affected Dhh1 expression levels, with mCh2-Dhh1 expression ∼50% lower than GFP-Dhh1 (Figure 3C and D). We can therefore not exclude that the reduction of P-bodies in the mCh2-Dhh1 strain is an effect of protein expression. However, this is not the case for His-mCherry-Dhh1, which has similar expression levels to GFP-Dhh1 yet displays a ∼40% reduction in P-body number.

**Figure 3.**
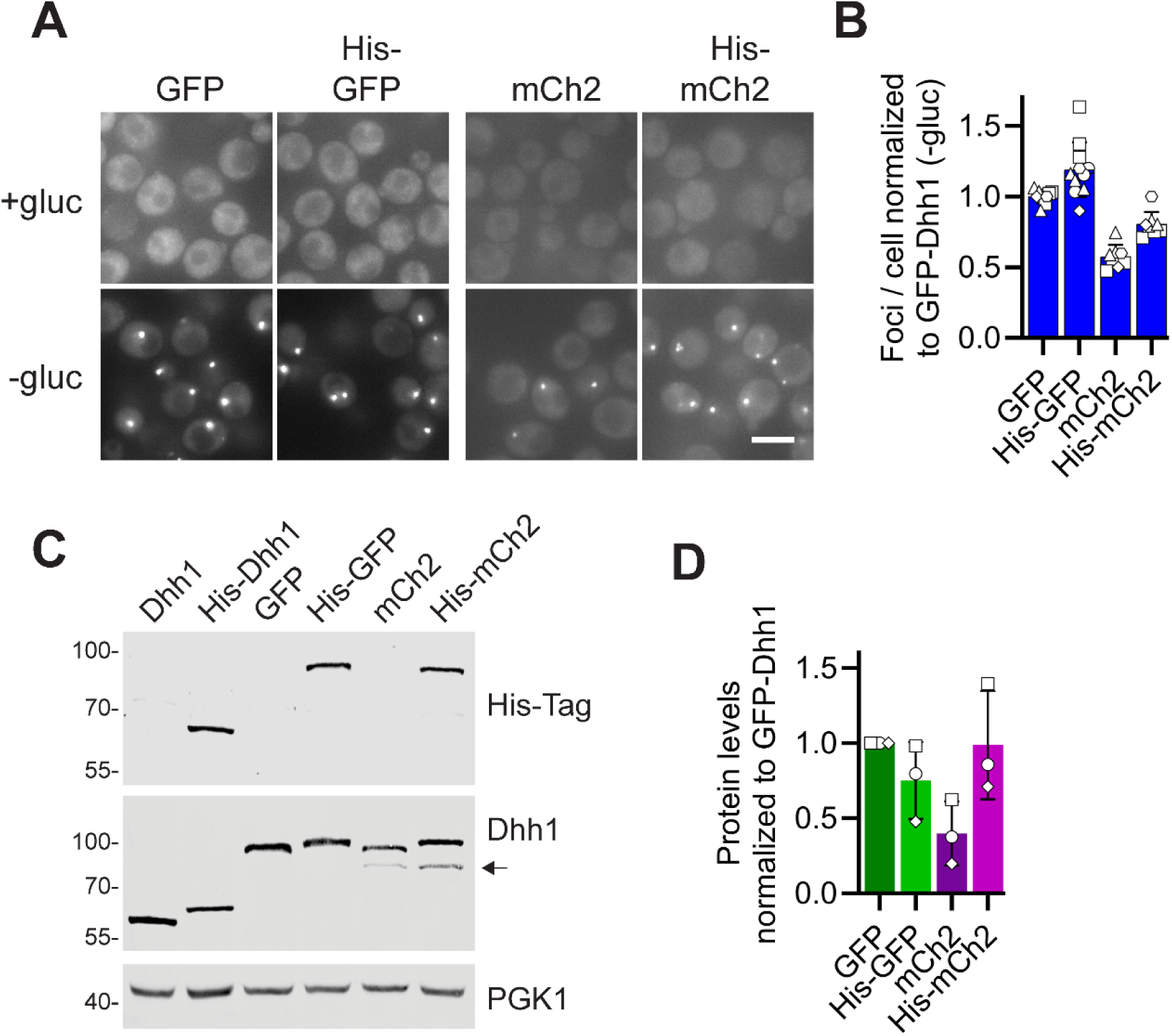
N-terminally tagged Dhh1 variants display variable number of P-bodies *in vivo*. (A) mCh2-Dhh1 shows different P-body numbers upon 30-minute glucose starvation compared to GFP-Dhh1. Scale bar = 5 μm. (B) Quantification of P-body numbers in (A) normalized to GFP-Dhh1 foci. N = 5, biological replicates are indicated with different symbols. (C) Western blot of logarithmically growing yeast cells expressing different Dhh1 constructs. Note that mCh2 tagging results in the appearance of a second band indicated by the arrow. (D) Quantification of (C) normalized to GFP-Dhh1 protein levels. N = 3, biological replicates are indicated with different symbols.

To get a wider overview of the effect of fluorescent tags on *in vivo* P-body formation, we fused mTurquoise2, mGFP, mNeonGreen, mVenus, mCherry, mKate2, and mRuby2 to Dhh1 at the endogenous locus, both N-and C-terminally. Western blotting showed that N-terminal tagging resulted in comparable expression levels for all Dhh1 constructs except the poorly expressed mCherry variant. By contrast, C-terminal tagging led to variable expression levels, with a ∼50% reduction for mNeonGreen, mCherry, mKate2 and mRuby2 compared to mTurquoise2 (Figure 4A-C).

**Figure 4.**
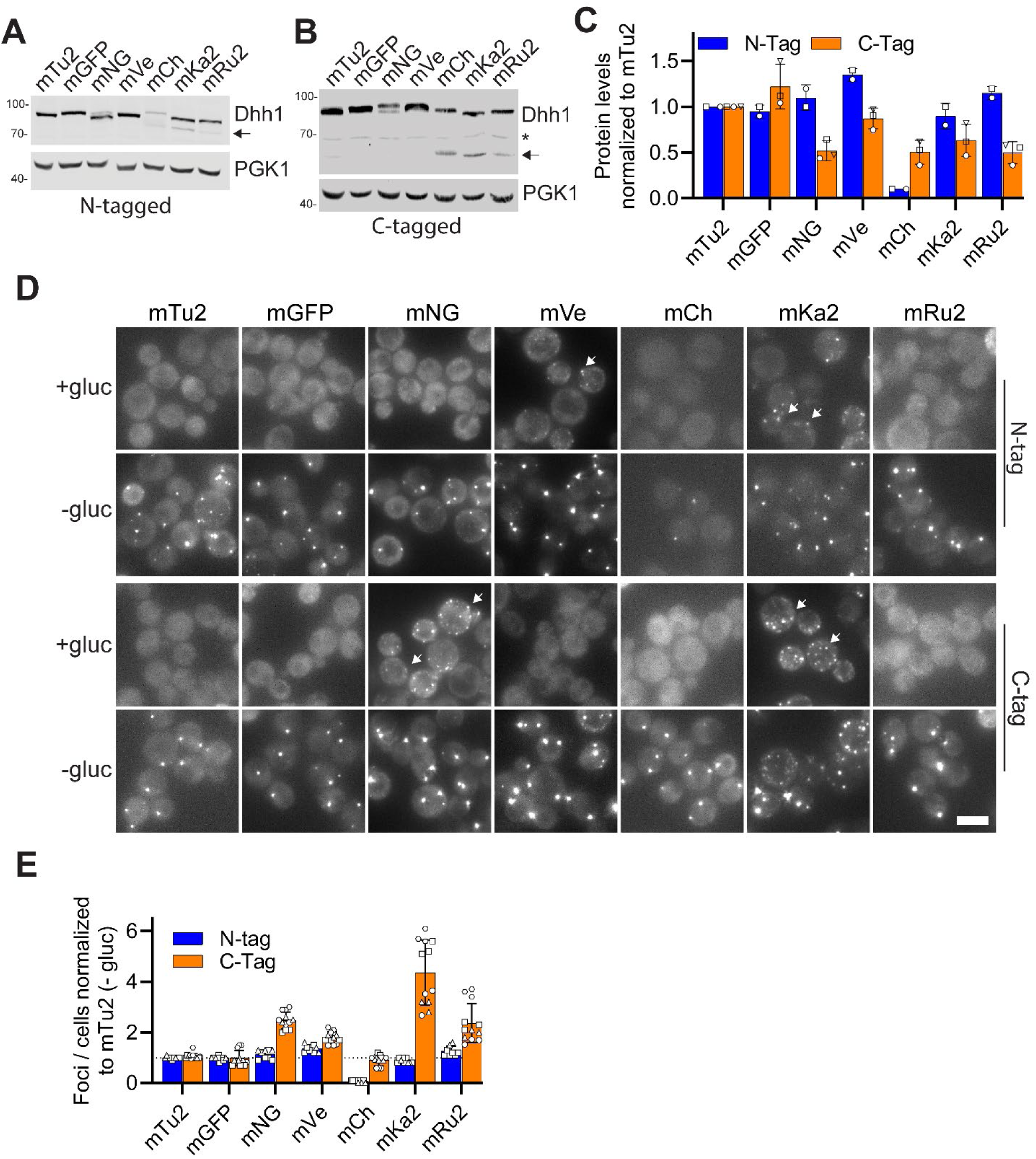
Position and type of fluorescent tag affect protein expression and P-body numbers *in vivo.* (A and B) Western blot of logarithmically growing yeast cells expressing N-tagged (A) or C-tagged (B) Dhh1. Note that mNeonGreen, mCherry, mKate2, and mRuby2 show an additional band indicated by the arrow. * indicates nonspecific band. (C) Quantification of (A) and (B) normalized to mTurquoise2-Dhh1 protein levels. N = 2 or 3, biological replicates are indicated with different symbols. (D) Position and type of fluorescent tag on endogenous Dhh1 affect P-body numbers upon 30-minute glucose starvation. Upper panel: N-tagged Dhh1. Lower panel: C-tagged Dhh1. The arrows indicate the presence of P-bodies in +glucose. Epifluorescent acquired images. Scale bar = 5 μm. (E) Quantification of the number of P-bodies per cell in (D). Values are normalized to mTurquoise2 foci. N = 4, biological replicates are shown with different symbols. mTu2, mTurquoise2; mGFP, mGFP; mNG, mNeonGreen; mVe, mVenus; mCh, mCherry; mKa2, mKate2; mRu2, mRuby2.

We then imaged and quantified the number of P-bodies per cells in +/-glucose using epifluorescence microscopy (Figure 4D and E) and confirmed the localization by spinning disk confocal microscopy (Supplemental Figure S4A). Upon glucose starvation, N-terminally tagged Dhh1 constructs had similar P-body numbers, except for mCherry-Dhh1, which showed none, consistent with its poor expression levels. In contrast, C-terminally tagged Dhh1 variants displayed striking differences upon glucose deprivation, with Dhh1-mKate2 showing the highest number of P-bodies (Figure 4D and E). Although mNeonGreen, mCherry, mKate2 and mRuby2 were expressed at similar levels, the number of P-bodies varied considerably between strains. This suggests that these changes in P-body numbers are not simply caused by variations in Dhh1 protein levels but instead are directly elicited by the addition of the tags.

Of note, some tagged constructs led to constitutive Dhh1 foci formation even in +gluc conditions (Dhh1-mNeonGreen, Dhh1-mKate2, mVenus-Dhh1 and mKate2-Dhh1; Figure 4D). For the C-terminally tagged constructs, we confirmed that these +gluc foci are *bona fide* P-bodies as they co-localized with Dcp2, another P-body marker (Supplemental Figure S4B). Intriguingly, Dcp2-GFP almost completely co-localized with Dhh1-mKate2 whereas Dcp2-mCherry showed much less co-localization with Dhh1-mNeonGreen in glucose rich medium, indicating that tagging of Dcp2 also affects its expression and/or localization in cells.

It is possible that the addition of tags, either through affecting Dhh1 expression levels or otherwise, would stress the cells and therefore induce P-body formation already in glucose-rich conditions. To examine this, we measured the growth of the Dhh1-tagged strains with wild-type (untagged Dhh1) and *dhh1Δ* strains as controls. Tagging of Dhh1 did not measurably affect the growth rate of any strain except N-terminally tagged mCherry-Dhh1, presumably due to its low expression levels. (Supplemental Figure S4C).

In sum, these data show that the tag itself and its position can greatly affect both protein expression and P-body number and can even lead to constitutive P-body formation in logarithmically growing, non-stressed cells.

### *In vivo* spike-in of Dhh1-GFP or Dhh1-mKate2 results in variations in P-body numbers

Given the different effects induced by the addition of tags, we attempted to find a method to reliably obtain a qualitative and quantitative readout for P-body formation *in vivo*. Similar to our *in vitro* experiments (Figure 2A-D), we sought to spike in a small amount of fluorescently tagged protein on top of untagged Dhh1 in cells. To this end, we expressed tagged copies of Dhh1 under the control of the low expression promoter *pREV1*, in addition to the endogenous untagged, wild type Dhh1 copy. We selected C-terminal GFP and mKate2 tags because they showed no and high P-body numbers in glucose-rich medium, respectively, and a ∼4 times difference in the number of P-bodies upon glucose deprivation (Figure 4E). Western blotting showed that expression of Dhh1-GFP and Dhh1-mKate2 under the *pREV1* promoter indeed led to very low expression compared to endogenous Dhh1 (Figure 5A), and we quantified that the fraction of tagged protein represented ∼2-4% of the total Dhh1 pool (Figure 5B). Surprisingly, even at these low protein levels the presence of tagged proteins affected P-body numbers both in the presence and absence of glucose (Figure 5C and D). Spiking-in of Dhh1-mKate2 led to ∼3 times more P-bodies compared to Dhh1-GFP and resulted in P-body formation even in the +gluc condition. Surprisingly, the presence of even small amounts of either Dhh1-GFP or Dhh1-mKate2 spike-ins led to the same number of P-bodies per cell in -gluc when compared to the expression of the respective, endogenously tagged Dhh1 variant as the only copy (Figure 5C, lower panel and Figure 5D). Together these results show that the presence of a tag can affect Dhh1 condensation and P-body formation *in vivo*, even when present only on a small fraction of proteins.

**Figure 5.**
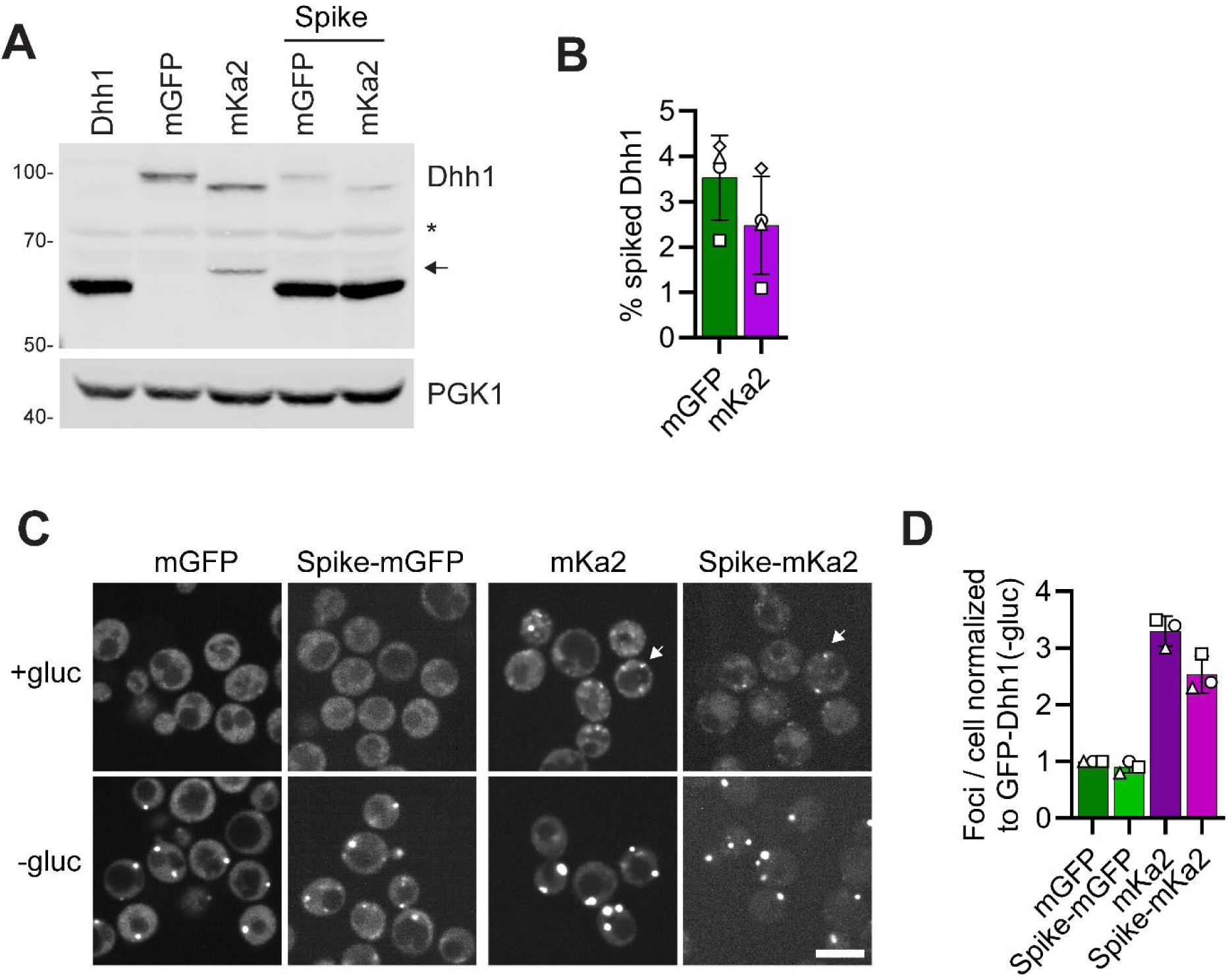
*In vivo* spiking-in of tagged Dhh1 affects P-body numbers. (A) Western blot of Dhh1-GFP and Dhh1-mKate2 either endogenously tagged or expressed from the p*Rev1* promoter integrated in the *TRP1* locus. The arrow indicates a cleaved product of Dhh1-mKate2. * indicates a nonspecific band. (B) Quantification of tagged Dhh1 relative to the endogenous Dhh1 protein levels. N = 4. (C) Spiking-in of Dhh1-GFP or Dhh1-mKate on top of an endogenously tagged Dhh1 variant does not further alter P-body numbers in +/-glucose compared to endogenously tagged Dhh1. Scale bar = 5 μm. (D) Quantification of the number of P-bodies in (C) normalized to GFP-Dhh1 P-bodies. N = 3, biological replicates are shown with different symbols.

In summary, the results presented here demonstrate that the addition of protein tags can drastically alter the condensation properties of Dhh1 *in vitro* and also affect P-body formation *in vivo*. We focus on Dhh1 in this study, yet our work raises a general concern about the use of protein tags to examine biomolecular condensation. Due to their dynamic nature and reliance on weak, multivalent interactions, biomolecular condensation reactions might be particularly sensitive to subtle changes in interaction strength or steric effects introduced by the addition of bulky protein tags that might impact packing density. Given our observations and additional findings by others that come to similar conclusions regarding the impact of protein tags on condensation (Martin *et al*., 2021; Zhou and Narlikar, 2023; Barkley *et al*., 2024; Dörner *et al*., 2024; Pandey *et al*., 2024), it might be prudent to re-examine some previously published results that exclusively relied on the use of tagged protein variants to study the formation and dynamics of biomolecular condensates and membraneless organelles. Going forward, it will be critical to carefully evaluate in each case the effects of protein tags on protein condensation reactions, both *in vitro* and *in vivo*.

Unfortunately, our results do not offer a simple solution to establish the ‘ground truth’. Spike-ins of a small fraction of tagged proteins might solve the problem in some instances, e.g., to study bulk condensation behavior. However, we were surprised to find that even very small fractions of tagged proteins can affect the number of P-bodies in cells (Figure 5) or the dynamics of condensates *in vitro* (Figure 2). In the latter case, this is likely due to the fact that only the tagged protein fraction can be examined, and the untagged protein is not visible in these assays. The effects of protein tags are particularly problematic for *in vivo* assays, where biomolecular condensates or membraneless organelles are not easily detectable without them and the use of antibodies requires fixation, preventing any dynamic measurements in living cells (Irgen-Gioro *et al*., 2022). In the future, it might therefore be important to establish label-free methods to examine condensate dynamics. The use of bio-orthogonal labeling methods that allow for the introduction of small fluorescent unnatural amino acids (Heil *et al*., 2018) might offer another alternative, since single small dyes might have no or only very small effects on condensation reactions.

We observed that the addition of some tags led to the formation of P-bodies even in unstressed, glucose-rich conditions, and intriguingly, this did not have any detectable impact on cell growth. Thus, the addition of protein tags might allow for functional experiments to address the physiological role of P-bodies, and it could now be interesting to carefully examine the impact of tagged Dhh1 variants on cell fitness and survival and on translation or mRNA turnover during stress. In general, protein tags might thus be useful tools to manipulate the condensation behavior of proteins without affecting their core functionality and could provide novel insight into the poorly understood physiological functions of biomolecular condensates in cells.

## Materials and Methods

### Yeast strain and plasmid construction

*Saccharomyces cerevisiae* strains were created in the W303 strain background using standard yeast genetic techniques including the transformation of a PCR product with homology regions to the yeast genome (Longtine *et al*., 1998). Plasmids were generated using restriction-based cloning methods or with the Golden Gate cloning system (Lee *et al*., 2015). For *in vivo* spike-in of Dhh1-GFP and Dhh1-mKate2, a linearized plasmid was integrated in the *TRP1* locus. For bacterial expression, Dhh1 constructs were cloned into modified pET-based vector with an N-terminal His-tag with a TEV cleavage site for removal of the tag (Bogomolovas *et al*., 2009; Fatti *et al*., 2023). Strains, plasmids and primers that were used in this study are listed in Supplemental Tables 1-3.

### Protein expression and purification

Dhh1 constructs were expressed as His-tagged proteins in *E. coli* Rosetta^TM^ (DE3) (Novagen) with an auto-induction medium system. Cells were grown at 37 °C in 1 L ZY complete medium to an OD_600_ of 0.7, then the temperature was lowered to 20 °C and growth continued for 20 h at 220 rpm. Cells were harvested and washed in cold PBS and the cell pellet flash frozen in liquid nitrogen before being stored at -20 °C for later use. The pellet was dissolved in 10 mL/g of Lysis Buffer (20 mM HEPES-KOH pH 7.7, 500 mM KCl, 5 mM MgCl_2_, 0.2 % IGEPAL CA-630, 10 mM Imidazole) supplemented with 0.5 mg/mL lysozyme (PanReac AppliChem) and 0.01 mg/mL DNAse I (PanReac AppliChem), and mechanically disrupted by high pressure homogenizer (Emulsiflex C5, Avestin). The lysate was centrifuged at 20’000 *g* (SS-34 fixed angle rotor, Sorvall). The supernatant was filtered through a 0.45 µm PES membrane filter (Sarstedt) and incubated with Ni-NTA beads (Qiagen). Beads were washed with 5 column volumes each of Detergent Buffer (20 mM HEPES-KOH pH 7.7, 500 mM KCl, 5 mM MgCl_2_, 0.2 % IGEPAL CA-630, 10 mM Imidazole, 5 % Glycerol), High Salt Buffer (20 mM HEPES-KOH pH 7.7, 1.5 M KCl, 5 mM MgCl_2_, 10 mM Imidazole, 5 % Glycerol), Imidazole Buffer (20 mM HEPES-KOH pH 7.7, 500 mM KCl, 5 mM MgCl_2_, 20 mM Imidazole, 5 % Glycerol), and eluted in 2.5 column volumes of Elution Buffer (20 mM HEPES-KOH pH 7.7, 500 mM KCl, 5 mM MgCl_2_, 330 mM Imidazole, 5 % Glycerol). The Elution Buffer was exchanged to Imidazole Buffer by passing the proteins through a PD-10 column (GE Healthcare) and for the His-cleaved samples incubated overnight at 10 °C with His-TEV protease. Eluate incubated with His-TEV was passed again through Ni-NTA beads to remove uncleaved proteins and His-TEV protease, concentrated with centrifugal filter units (Millipore) and further purified by size exclusion chromatography on a Superdex 200 16/600 column (GE Healthcare) in Storage Buffer (30 mM HEPES-KOH pH 7.7, 500 mM KCl, 5 mM MgCl_2_, 1 mM DTT, 10 % Glycerol) using an AKTA pure system (GE Healthcare). Proteins not incubated with His-TEV protease were immediately processed with size exclusion chromatography. Positive fractions were pooled, further concentrated and final purity assessed by SDS-PAGE and Coomassie stain (Instant Blue®, Abcam). 10 μL aliquots were flash frozen in PCR tubes and stored at -80 °C.

### N-labeling of Dhh1

Dhh1 was chemically labeled with Cy5-NHS-ester dye (Cyanine 5 SE, Tocris). Labeling reactions were performed at pH 6.0. Dhh1 and Cy5-NHS-ester dye were mixed in a molar ratio of 1:4, followed by an incubation of 1 h at room temperature, protected from light. Free dye was twice removed with PD-10 columns. Cy5-Dhh1 was aliquoted, snap-frozen in liquid nitrogen and stored at -80 °C. Protein concentration and average degree of labeling was determined following the manufacturer’s instructions.

### Western blot

Roughly 5 OD_600_ units of log growing cells were harvested and treated with 0.2 M NaOH for 15 minutes at room temperature, then centrifuged and the NaOH supernatant removed. The cell pellet was dissolved in SDS sample buffer and boiled at 95 °C for 6 minutes. Proteins were resolved by 10 % or 12 % Tris-Glycine SDS-PAGE (Biorad), then transferred to nitrocellulose membrane (GE Life Sciences, Marlborough, MA) with a semi-dry transfer using Trans-Blot® Turbo™ (BioRad). Membranes were blocked in PBS with 5 % non-fat milk, followed by incubation with the primary antibody overnight. Membranes were washed four times with PBS with 0.05 % Tween-20 (PBS-T) and incubated with a secondary antibody for 1 h at room temperature. Membranes were imaged and protein bands quantified using an infrared imaging system (Odyssey; LI-COR Biosciences, Lincoln, NE). All primary antibodies were diluted in 5% BSA in PBS-T and secondary antibodies in 5 % non-fat milk in PBS-T. Mouse anti-His tag 1:2’000, Rabbit anti-Dhh1 1:8’000, Mouse anti-PGK1 (22C5D8) 1:5’000, IRDye® 800CW Goat anti-Rabbit IgG (H + L) 1:25’000, Alexa 680 Goat anti-Mouse 1:25’000.

### *In vitro* droplet assay

Dhh1 protein stock was diluted in Storage Buffer to an intermediate dilution to obtain 100 mM KCl, except for the KCl titration assay where the minimal KCl concentration was 50 mM. This passage was required to keep constant the amount of salt and other components brought in by the Storage Buffer. 100 mM ATP stock solution was neutralized to pH 7.4 by addition of 10 mM HEPES-KOH and 1N NaOH and always complexed with MgCl_2_. PolyUridilic acid (pU, Sigma) was dissolved in water, stored in aliquots at 1 or 10 mg/mL and further diluted to the required final amount. The final buffer condition for each assay is the following without the titrated component: 30 mM MES-KOH pH 6.0, 100 mM KCl, 50 ng/μL pU, 2 mM MgCl_2_, 5 mM ATP/MgCl_2_, 0.5 mM DTT. For the pH titration assay, 30 mM MES-KOH was used for pH 5.8-6.2, and PIPES-KOH for pH 6.4-7.4. The reaction was directly performed in a 384-well glass-bottom plate in a final volume of 20 μL. Proteins solutions were kept separated from pU and at low pH to avoid condensation in the tube before pipetting. Condensates were imaged with a Nikon Eclipse Ti Microscope with a 60X objective after at least 30 minutes to allow the condensates to settle at the bottom of the plate. For the spiking in assay, GFP-Dhh1, mCh2-Dhh1, and Cy5-Dhh1 protein stocks were adjusted to give the final required KCl concentration when mixed with untagged Dhh1 and processed as described.

### Microscopy

Yeast cells were grown overnight in complete minimal synthetic media (SCD), diluted to OD_600_ 0.2 the following day, and grown to mid-log phase (OD_600_ ∼ 0.6). 20 μL cells were directly transferred onto a Concanavalin A-treated 384-well plate (Brooks, Matriplate) and washed 3 times with 80 μL of either SCD or SC, centrifuged down at 100 g every wash, and finally 50 μL were added before imaging. Cells were incubated for at least 30 minutes before imaging and the incubation time was recorded after the first wash. Cells were imaged either with epifluorescence or spinning disk microscopy.

Epifluorescence images were acquired with a Nikon Ti 7 Eclipse equipped with a Spectra X LED light source (Lumencore) using an Apochromat VC 100x objective NA 1.4 (Nikon) (filters: Spectra emission filters 475/28 & 542/27 and DAPI/FITC/Cy3/Cy5 Quad HC Filterset with 410/504/582/669 HC Quad dichroic and a 440/521/607/700 HC quadband filter (Semrock)) equipped with the NIS Elements software (Nikon). Images were acquired as z-stacks with a Flash 4.0 sCMOS camera with 0.5 um sections (Hamamatsu) and processed using ImageJ software with an exposure time of 100-400 ms.

For spinning disk imaging, a single plane of yeast cells was acquired on an inverted Nikon spinning disk microscope equipped with the Yokogawa Confocal Scanner Unit CSU-W1-T2 SoRa and a triggered Piezo z-stage (Mad City Labs Nano-Drive). It was used in spinning disk mode with a pinhole diameter of 50 μm combined with a 1.45 NA, 100x objective and controlled by the NIS Elements Software (Nikon). Images were acquired as single stacks with a sCMOS Hamamatsu Orca Fusion BT camera (2304 x 2304 pixel, 6.5 x 6.5 μm pixel size). Imaging was performed at RT with 80 % laser intensity of a DPSS 488 nm 200 mW and/or Diode 561 nm, 200 mW light source with exposure times of 100-200 ms.

For yeast P-body quantification we generated a maximum projection of xz central z-slices in FIJI (Schindelin *et al*., 2012). For cell segmentation we used a custom script written in Jython (FIJI) (Schindelin *et al*., 2012) that calls and applies the CellPose (Stringer *et al*., 2021) framework in batch to our fluorescent images. We used the pretrained model Cyto2 with an initial diameter of 4 μm. The number of foci was calculated using the FIJI (Schindelin *et al*., 2012) plugin Trackmate (Ershov *et al*., 2022). using the LoG detector method with a radius of 400 nm and a variable quality threshold depending on the fluorescent protein.

### Fluorescence Recovery After Photobleaching (FRAP)

Fluorescence recovery after photobleaching (FRAP) assays of *in vitro* condensates were set up in a 384-well plate. The final assay conditions were: 30 mM MES-KOH pH 6.0-7.0, 100 mM KCl, 50 ng/μL pU, 2 mM MgCl_2_, 5 mM ATP/MgCl_2_, 0.5 mM DTT. Condensates were bleached by focusing a 488 nm or 561 nm laser light on a circular area with a diameter of ∼ 2 μm. Image analysis, including background subtraction, correction of bleaching during recovery, and normalization to pre-and postbleach intensity was performed via the webtool easyFRAP-web (https://easyfrap.vmnet.upatras.gr/) (Koulouras *et al*., 2018) and further analysis was performed with GraphPad Prism 10.

### Growth curve

Overnight yeast cultures were grown in permissive conditions in SCD media. Growth curves were acquired using CLARIOstar automated plate reader (BMG Labtech, Ortenberg, Germany) at 30 °C in 48-well plastic plates in duplicate (Thermo Fisher) from overnight pre-cultures diluted to OD_600_ 0.15 in the specified synthetic liquid media.

## Abbreviations

pU: PolyUridilic acid
ATP: Adenosine Triphosphate
KCl: Potassium Chloride
His-tag: Histidine tag
Cy5: Cyanine 5
FRAP: Fluorescence Recovery After Photobleaching
LCDs: Low Complexity Domains

## Author contributions

Conceptualization: EF and KW. Methodology: EF and KW. Investigation: EF and KW. Formal analysis: EF. Writing – Original Draft: EF, SK and KW. Writing – Review & Editing: EF, SK and KW, Funding acquisition: KW. Supervision: KW.

## Acknowledgements

The authors are grateful to members of the Weis lab for discussions and suggestions on the manuscript. We would like to acknowledge Sachin Yuvaraj Dinesh Babu for assistance during the early stages of the manuscript. We thank ScopeM and Joachim Hehl for technical assistance and help with microscopy and Pablo Aurelio Gómez García for the custom script for cell segmentation and P-body quantification. This work was supported by grants from the Swiss National Science Foundation (SNF CRSII5_193740, 310030_208213 and TMAG-3_209354 to K.W.).

---

**Supplemental Figure S1.**
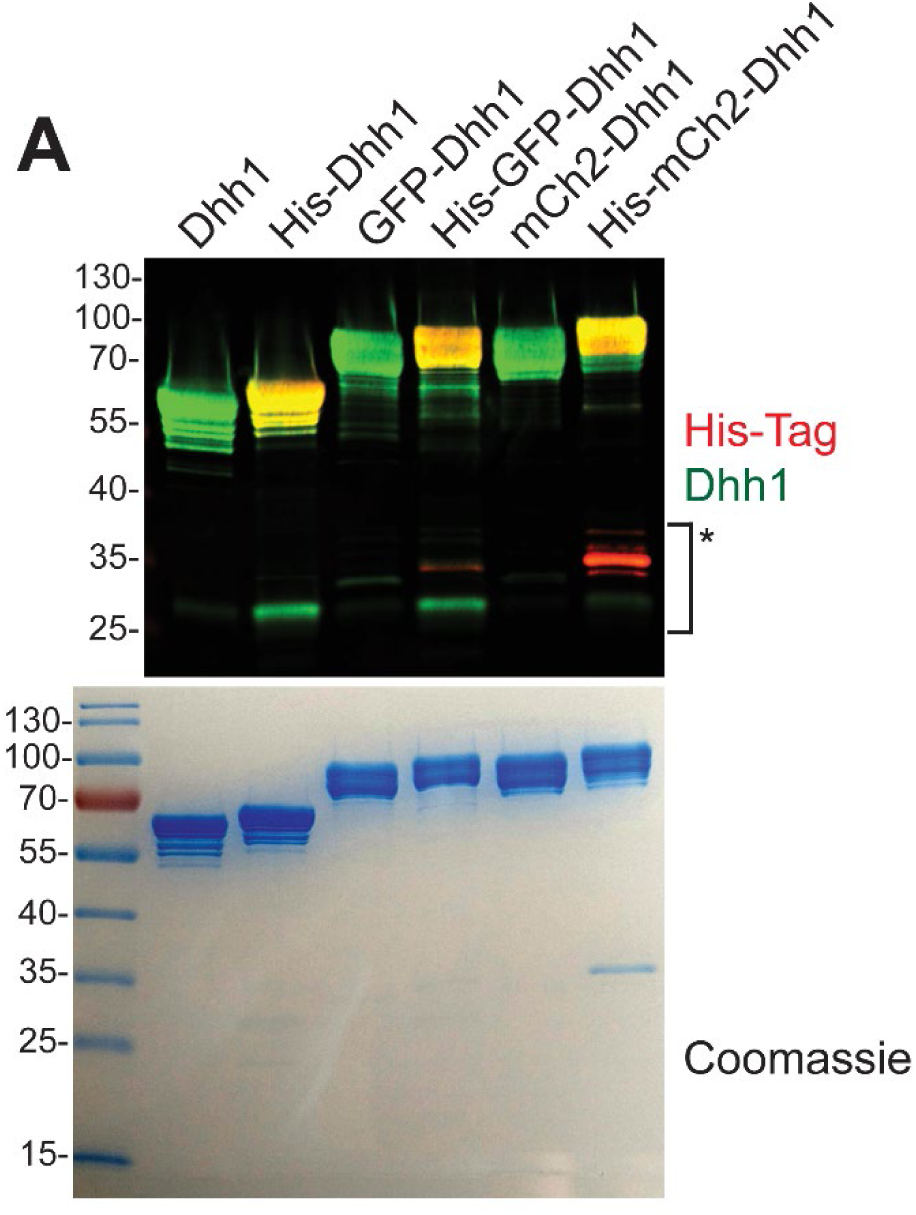
Dhh1, GFP-Dhh1 and mCh2-Dhh1 with or without a His tag can be expressed and purified from *E. coli*. (A) Western blot (upper) and Coomassie staining (lower panel) of the purified Dhh1 constructs used in this study. * indicates degradation products.

**Supplemental Figure S2.**
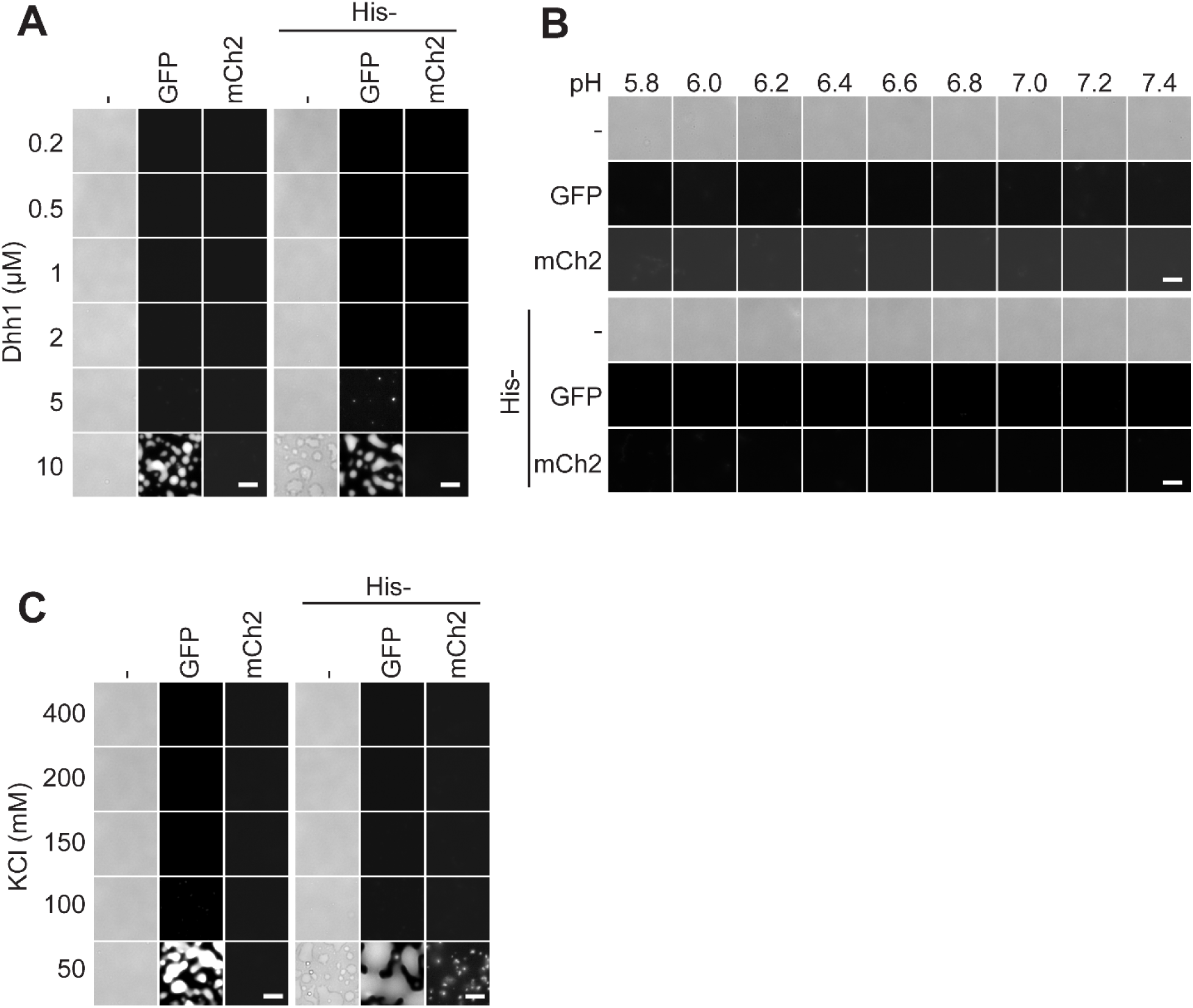
Tagged Dhh1 can form condensates without pU in certain conditions (A) Left panel: GFP-Dhh1 forms condensates at high concentrations. Right panel: His-Dhh1 and His-GFP-Dhh1 but not His-mCh2-Dhh1 form condensates at high concentrations. Assay performed in the absence of pU at pH 6.0. (B) At 5 μM, none of the Dhh1 constructs form condensates in the absence of pU at any pH tested. (C) Left panel: GFP-Dhh1 forms condensates at low salt concentrations. Right panel: His-Dhh1, His-GFP-Dhh1 and His-mCh2-Dhh1 form condensates at low salt concentrations. Assay performed in the absence of pU at pH 6.0. 5 mM ATP is used in all assays. Scale bar = 10 μm.

**Supplemental Figure S3.**
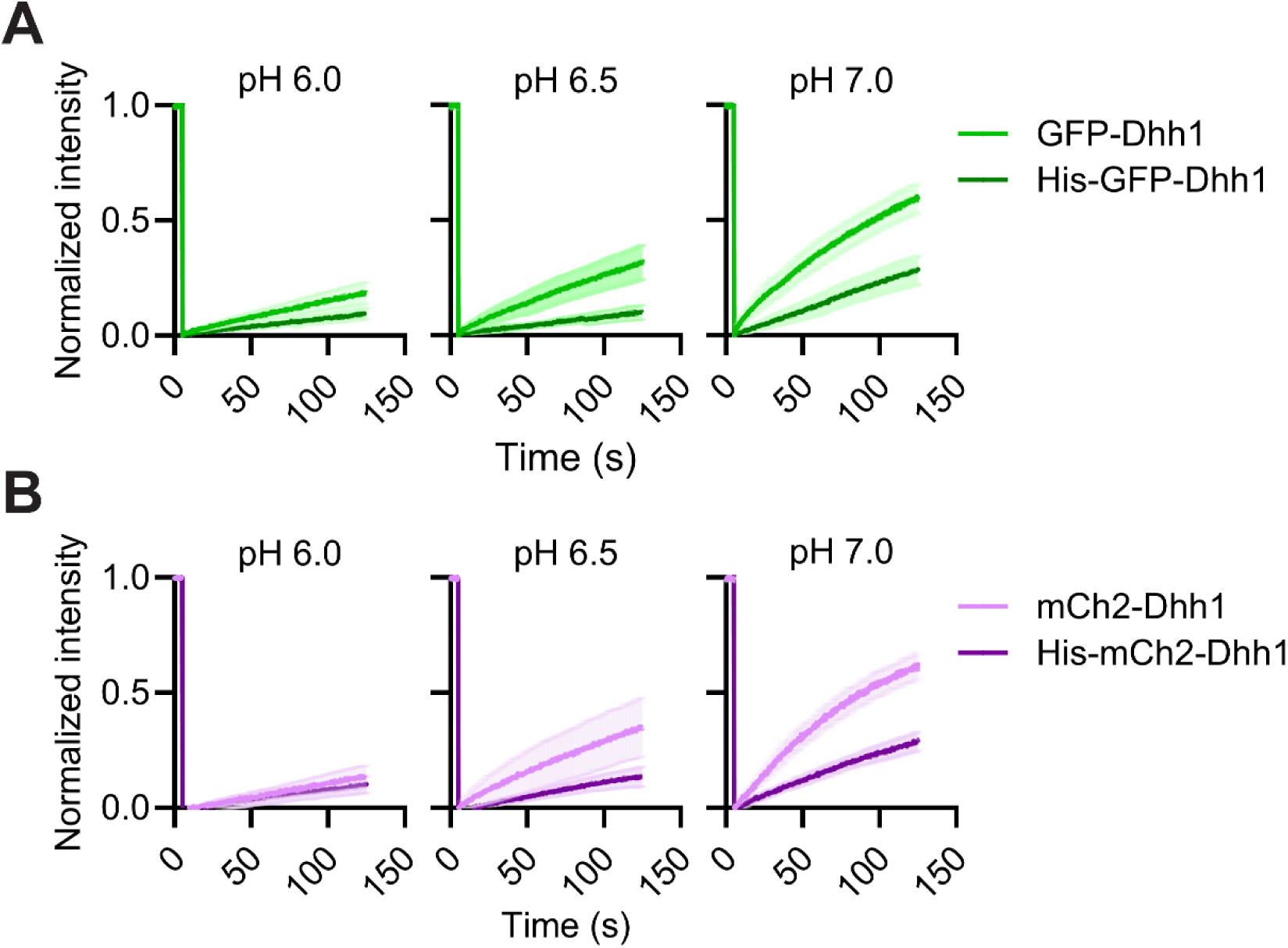
A His tag decreases FRAP recovery rates of doubly tagged Dhh1 constructs. (A and B) Spiking-in of 2 % GFP-Dhh1 and His-GFP-Dhh1 (A) or mCh2-Dhh1 and His-mCh2-Dhh1 (B) to 5 μM Dhh1 in the presence of pU at different pHs shows a consistently lower recovery rate for His-tagged Dhh1. N = 2.

**Supplemental Figure S4.**
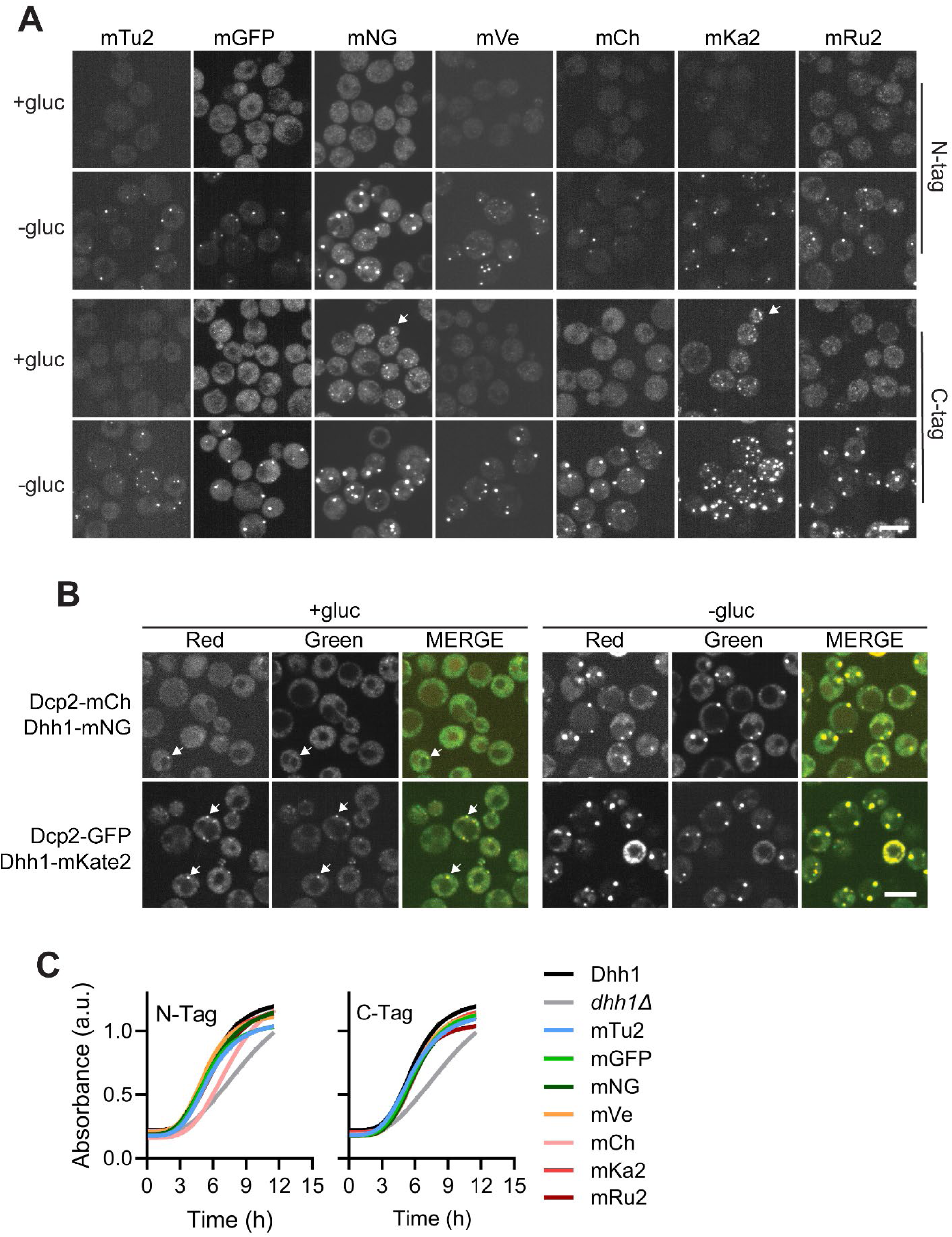
Dhh1 foci observed in non-stressed cells colocalize with P-body markers and variable P-body numbers do not seem to affect cell growth. (A) Spinning disk microscopy of single plane yeast strains expressing tagged Dhh1 in +/-glucose. (B) Dhh1-mNeonGreen or Dhh1-mKate2 colocalize with Dcp2 in +/-glucose. (C) Dhh1 tagging does not affect cell growth when Dhh1 is expressed at levels similar to the endogenous protein. Scale bar = 5 μm.

## References

Aguzzi, A, and Altmeyer, M (2016). Phase Separation: Linking Cellular Compartmentalization to Disease. Trends in Cell Biology 26, 547–558.

Aizer, A, Kalo, A, Kafri, P, Shraga, A, Ben-Yishay, R, Jacob, A, Kinor, N, and Shav-Tal, Y (2014). Quantifying mRNA targeting to P-bodies in living human cells reveals their dual role in mRNA decay and storage. Journal of Cell Science 127, 4443–4456.

Alberti, S, and Dormann, D (2019). Liquid-Liquid Phase Separation in Disease. Annu Rev Genet 53, 171–194.

Banani, SF, Lee, HO, Hyman, AA, and Rosen, MK (2017). Biomolecular condensates: organizers of cellular biochemistry. Nature Reviews Molecular Cell Biology 18, 285–298.

Barkley, RJR, Crowley, JC, Brodrick, AJ, Zipfel, WR, and Parker, JSL (2024). Fluorescent protein tags affect the condensation properties of a phase-separating viral protein. MBoC 35, ar100.

Bogomolovas, J, Simon, B, Sattler, M, and Stier, G (2009). Screening of fusion partners for high yield expression and purification of bioactive viscotoxins. Protein Expr Purif 64, 16–23.

Carroll, JS, Munchel, SE, and Weis, K (2011). The DExD/H box ATPase Dhh1 functions in translational repression, mRNA decay, and processing body dynamics. J Cell Biol 194, 527–537.

Choi, J-M, Holehouse, AS, and Pappu, RV (2020). Physical Principles Underlying the Complex Biology of Intracellular Phase Transitions. Annual Review of Biophysics 49, 107–133.

Cranfill, PJ, Sell, BR, Baird, MA, Allen, JR, Lavagnino, Z, de Gruiter, HM, Kremers, G-J, Davidson, MW, Ustione, A, and Piston, DW (2016). Quantitative assessment of fluorescent proteins. Nat Methods 13, 557–562.

Derrer, CP, Mancini, R, Vallotton, P, Huet, S, Weis, K, and Dultz, E (2019). The RNA export factor Mex67 functions as a mobile nucleoporin. Journal of Cell Biology 218, 3967–3976.

Dörner, K, Gut, M, Overwijn, D, Cao, F, Siketanc, M, Heinrich, S, Beuret, N, Sharpe, T, Lindorff-Larsen, K, and Maria, H (2024). Tag with Caution — How protein tagging influences the formation of condensates. 2024.10.04.616694.

Ershov, D, Phan, M-S, Pylvänäinen, JW, Rigaud, SU, Le Blanc, L, Charles-Orszag, A, Conway, JRW, Laine, RF, Roy, NH, Bonazzi, D, et al. (2022). TrackMate 7: integrating state-of-the-art segmentation algorithms into tracking pipelines. Nat Methods 19, 829–832.

Fatti, E, Hirth, A, Švorinić, A, Günther, M, Stier, G, Cruciat, C-M, Acebrón, SP, Papageorgiou, D, Sinning, I, Krijgsveld, J, et al. (2023). DEAD box RNA helicases act as nucleotide exchange factors for casein kinase 2. Science Signaling 16, eabp8923.

Feric, M, Vaidya, N, Harmon, TS, Mitrea, DM, Zhu, L, Richardson, TM, Kriwacki, RW, Pappu, RV, and Brangwynne, CP (2016). Coexisting Liquid Phases Underlie Nucleolar Subcompartments. Cell 165, 1686–1697.

Heil, CS, Rittner, A, Goebel, B, Beyer, D, and Grininger, M (2018). Site-Specific Labelling of Multidomain Proteins by Amber Codon Suppression. Sci Rep 8, 14864.

Hirose, T, Ninomiya, K, Nakagawa, S, and Yamazaki, T (2023). A guide to membraneless organelles and their various roles in gene regulation. Nat Rev Mol Cell Biol 24, 288–304.

Hondele, M, Heinrich, S, De Los Rios, P, and Weis, K (2020). Membraneless organelles: phasing out of equilibrium. Emerg Top Life Sci 4, 331–342.

Hondele, M, Sachdev, R, Heinrich, S, Wang, J, Vallotton, P, Fontoura, BMA, and Weis, K (2019). DEAD-box ATPases are global regulators of phase-separated organelles. Nature 573, 144–148.

Hyman, AA, Weber, CA, and Jülicher, F (2014). Liquid-Liquid Phase Separation in Biology. Annual Review of Cell and Developmental Biology 30, 39–58.

Irgen-Gioro, S, Yoshida, S, Walling, V, and Chong, S (2022). Fixation can change the appearance of phase separation in living cells. ELife 11, e79903.

Koulouras, G, Panagopoulos, A, Rapsomaniki, MA, Giakoumakis, NN, Taraviras, S, and Lygerou, Z (2018). EasyFRAP-web: a web-based tool for the analysis of fluorescence recovery after photobleaching data. Nucleic Acids Research 46, W467–W472.

Lambert, TJ (2019). FPbase: a community-editable fluorescent protein database. Nat Methods 16, 277–278.

Lee, ME, DeLoache, WC, Cervantes, B, and Dueber, JE (2015). A Highly Characterized Yeast Toolkit for Modular, Multipart Assembly. ACS Synth Biol 4, 975–986.

Longtine, MS, McKenzie, A, Demarini, DJ, Shah, NG, Wach, A, Brachat, A, Philippsen, P, and Pringle, JR (1998). Additional modules for versatile and economical PCR-based gene deletion and modification in Saccharomyces cerevisiae. Yeast 14, 953–961.

Luo, Y, Na, Z, and Slavoff, SA (2018). P-Bodies: Composition, Properties, and Functions. Biochemistry 57, 2424–2431.

Martin, EW, Thomasen, FE, Milkovic, NM, Cuneo, MJ, Grace, CR, Nourse, A, Lindorff-Larsen, K, and Mittag, T (2021). Interplay of folded domains and the disordered low-complexity domain in mediating hnRNPA1 phase separation. Nucleic Acids Research 49, 2931–2945.

Mugler, CF, Hondele, M, Heinrich, S, Sachdev, R, Vallotton, P, Koek, AY, Chan, LY, and Weis, K (2016). ATPase activity of the DEAD-box protein Dhh1 controls processing body formation. ELife 5, e18746.

Pandey, NK, Varkey, J, Ajayan, A, George, G, Chen, J, and Langen, R (2024). Fluorescent protein tagging promotes phase separation and alters the aggregation pathway of huntingtin exon-1. Journal of Biological Chemistry 300.

Parker, R, and Sheth, U (2007). P Bodies and the Control of mRNA Translation and Degradation. Molecular Cell 25, 635–646.

Roden, C, and Gladfelter, AS (2021). RNA contributions to the form and function of biomolecular condensates. Nat Rev Mol Cell Biol 22, 183–195.

Schindelin, J, Arganda-Carreras, I, Frise, E, Kaynig, V, Longair, M, Pietzsch, T, Preibisch, S, Rueden, C, Saalfeld, S, Schmid, B, et al. (2012). Fiji: an open-source platform for biological-image analysis. Nat Methods 9, 676–682.

Schlüßler, R, Kim, K, Nötzel, M, Taubenberger, A, Abuhattum, S, Beck, T, Müller, P, Maharana, S, Cojoc, G, Girardo, S, et al. (2022). Correlative all-optical quantification of mass density and mechanics of subcellular compartments with fluorescence specificity. ELife 11, e68490.

Sheth, U, and Parker, R (2003). Decapping and decay of messenger RNA occur in cytoplasmic processing bodies. Science 300, 805–808.

Shin, Y, and Brangwynne, CP (2017). Liquid phase condensation in cell physiology and disease. Science 357, eaaf4382.

Snapp, EL (2009). Fluorescent proteins: a cell biologist’s user guide. Trends in Cell Biology 19, 649–655.

Stringer, C, Wang, T, Michaelos, M, and Pachitariu, M (2021). Cellpose: a generalist algorithm for cellular segmentation. Nat Methods 18, 100–106.

Teixeira, D, and Parker, R (2007). Analysis of P-body assembly in Saccharomyces cerevisiae. Molecular Biology of the Cell 18, 2274–2287.

Uebel, CJ, and Phillips, CM (2019). Phase-separated protein dynamics are affected by fluorescent tag choice. MicroPublication Biology.

Zhou, Z (Kate), and Narlikar, GJ (2023). Understanding how genetically encoded tags affect phase separation by Heterochromatin Protein HP1α, Biophysics.

