## Supplemental Table for "The dark side of fluorescent protein tagging – the impact of protein tags on biomolecular condensation"

| Plasmid name (GGKW) | Plasmid name | Golden Gate Part Type | Gene | Reference |
|---|---|---|---|---|
| GGKW1 | EntryVector | EMTPY | Empty | https://doi.org/10.1021/sb500366v |
| GGKW117 | 4a_mCherry | 4A | mCherry tag | this study |
| GGKW119 | 3a_mCherry | 3A | mCherry tag | this study |
| GGKW129 | 6_TRP1 | 6 | TRP1 marker. Consists of the TRP1 promotor, ORF and terminator | this study |
| GGKW142 | 2_PringleUptag(F1/F2) | 2 | Pringle Uptags: ccagctgaagcttcgtacgctgcaggtcgacggatccccgggttaattaa | this study |
| GGKW144 | 7_PringleDowntag(R1/R3/pkw1964) | 7 | Pringle Downtags: CAGTATAGCGACCAGCATTCaatacgcaaaccgcctctgaattcgagctcgtttaaac | this study |
| GGKW172 | 4a_yeGFP | 4A | Monomeric yEGFP(A206K) | this study |
| GGKW173 | 4a_mNeongreen | 4A | mNeongreen | this study |
| GGKW174 | 4a_mKate2 | 4A | mKate2. Nucleotide sequence was codon optimized for yeast expression | this study |
| GGKW175 | 3a_yeGFP | 3A | Monomeric yEGFP(A206K) | this study |
| GGKW176 | 3a_mNeongreen | 3A | mNeongreen | this study |
| GGKW177 | 3a_mKate2 | 3A | mKate2. Nucleotide sequence was codon optimized for yeast expression | this study |
| GGKW183 | 8b_Trp1 5' Homology | 8B | TRP1 5' homology region | this study |
| GGKW184 | 7_Trp1 3' Homology | 7 | TRP1 3' homology region | this study |
| GGKW2 | 1_ConLS | 1 | Default connector sequence left | https://doi.org/10.1021/sb500366v |
| GGKW222 | 3_Dhh1 | 3 | Dhh1 entry vector for C tagging | this study |
| GGKW231 | 3b_Dhh1 | 3B | Dhh1 entry vector for N tagging | this study |
| GGKW27 | 2_pREV1 | 2 | Promoter, Low strenght | https://doi.org/10.1021/sb500366v |
| GGKW37 | 3a_mTurquoise2 | 3A | mTurquoise2 | https://doi.org/10.1021/sb500366v |
| GGKW38 | 3a_Venus | 3A | mVenus | https://doi.org/10.1021/sb500366v |
| GGKW39 | 3a_mRuby2 | 3A | mRuby2 | https://doi.org/10.1021/sb500366v |
| GGKW51 | 4_tENO1 | 4 | Terminator | https://doi.org/10.1021/sb500366v |
| GGKW53 | 4_tADH1 | 4 | Terminator | https://doi.org/10.1021/sb500366v |
| GGKW57 | 4a_mTurquoise2 | 4A | mTurquoise2 | https://doi.org/10.1021/sb500366v |
| GGKW58 | 4a_Venus | 4A | mVenus | https://doi.org/10.1021/sb500366v |
| GGKW59 | 4a_mRuby2 | 4A | mRuby2 | https://doi.org/10.1021/sb500366v |
| GGKW63 | 4b_tADH1 | 4B | Terminator | https://doi.org/10.1021/sb500366v |
| GGKW67 | 5_ConR1 | 5 | Default Connector sequence right | https://doi.org/10.1021/sb500366v |
| GGKW76 | 6_HIS3 | 6 | HIS3 (ScHIS3 Promotor, HIS3, ScHIS3 Terminator) | https://doi.org/10.1021/sb500366v |
| GGKW83 | 8_AmpR-ColE1 | 8 | E. coli drug resistance marker for Ampicillin (default) and ColE1 origin | https://doi.org/10.1021/sb500366v |
| GGKW89 | 8a_AmpR-ColE1 | 8A | E. coli drug resistance marker for Ampicillin (default) and ColE1 origin | https://doi.org/10.1021/sb500366v |
